# Interleukin-22 mediated renal metabolic reprogramming via PFKFB3 to treat kidney injury

**DOI:** 10.1101/2020.08.04.237347

**Authors:** Wei Chen, Yilan Shen, Jiajun Fan, Xian Zeng, Xuyao Zhang, Jingyun Luan, Jinghui Zhang, Si Fang, Xiaobin Mei, Dianwen Ju

## Abstract

Kidney damage initiates the deteriorating metabolic states in tubule cells that lead to the development of end-stage renal disease (ESTD). Interleukin 22 (IL-22) is an effective therapeutic antidote for kidney injury via promoting kidney recovery, but little is known about the underlying molecular mechanisms. Here we first provide evidence that IL-22 attenuates kidney injury via metabolic reprogramming of renal tubular epithelial cells (TECs). Specifically, our data suggest that IL-22 regulates mitochondrial function and glycolysis in damaged TECs. Further observations indicate that IL-22 alleviates the accumulation of mitochondrial reactive oxygen species (ROS) and dysfunctional mitochondria via the induction of AMPK/AKT signaling and PFBFK3 activities. In mice, amelioration of kidney injury and necrosis and improvement of kidney functions via regulation of these metabolism relevant signaling and mitochondrial fitness of recombinant IL-22 are certificated in cisplatin induced kidney damage and diabetic nephropathy (DN) animal models. Taken together, our findings unravel new mechanistic insights into protective effects of IL-22 on kidney and highlight the therapeutic opportunities of IL-22 and the involved metabolic regulators in various kidney diseases.

## 1. Introduction

Acute kidney injury (AKI), a common public health concern associated with high mortality and morbidity, affects millions of hospitalized patients worldwide and shows a fast-increasing incidence [1-3]. The underlying pathophysiology of AKI is not fully understood but involves the damage and apoptosis of renal tubular epithelial cells (TECs), especially in the proximal tubule [4-7]. Multiple literatures support that their severe and sustained damage often causes incomplete and maladaptive tissue repair, leading to inflammatory response, tubular degeneration, kidney fibrosis and eventually progression to end-stage renal disease (ESTD) [8-10]. Unfortunately, other than dialysis-based supportive care, no well-established therapeutics for kidney injury are available. Therefore, proactive treatment is much needed to reduce the suffering it causes to patients and the significant financial burdens of kidney injury to individuals and society.

Interleukin-22 (IL-22), an important member of the IL-10 family cytokines, has recently attracted tremendous attention as a survival agent in numerous disorders driven by epithelial damage [11-13]. IL-22 elicits tissue protection and homeostasis primarily via activation of STAT3 signaling pathway and the promotion of epithelial proliferation [14-15]. Importantly, studies have demonstrated that the expression of IL-22R1 is limited to renal proximal TECs, and treatment with IL-22 can protect against renal epithelial injury and accelerate tubular regeneration [16-17]. However, although accumulating evidence indicates IL-22 is an effective therapeutic antidote for kidney injury, little is known about the underlying mechanisms of IL-22 induced TECs recovery. Understanding the molecular basis of IL-22 is significant both for exploring how IL-22 acts to inhibit AKI or ESTD and for discovering molecular modifiers as well as crucial processes involved in preventing kidney damage.

Kidney repair after damage is a metabolically-dependent and complicated process [18]. Recent evidence indicates that cell metabolic signatures can regulate renal cell survival and plasticity and thus provide useful mechanistic insights into how the metabolic processes control tissue repair [19-20]. Specifically, these findings show that metabolic reprogramming by PGC1α and the S-nitroso-CoA reductase system protects against kidney injury. Mechanistically, increased glycolysis and oxidative phosphorylation (OXPHOS) can promote growth and defensive signaling pathways or can supply the higher energetic demands of mitosis and anabolic biosynthesis during organ repair [19-21]. Recently, our studies have demonstrated that IL-22 alleviates mitochondrial dysfunction in liver, which involves the cellular metabolic processes [22-24]. Thus, we asked whether IL-22 protects against TECs damage and apoptosis via regulating their metabolic states. To determine the mechanisms of IL-22 mediated kidney repair, we treated human proximal tubular epithelial cells (PTECs) with stress stimulation. We found that IL-22 drives a metabolic reprogramming to enhance glycolysis and OXPHOS, which might prevent against TECs dysfunction. We further demonstrated that IL-22 induced this program via activating of AMPK/AKT signaling and its molecular modifiers to alleviate mitochondrial dysfunction. Our observations shed light on regulating metabolic states for treating and preventing kidney diseases, and provide new insights for developing efficient novel interventions for organ injury in general.

## 2. Results

### IL-22 preserves cellular metabolism in TECs

Because mitochondria have been suggested to be important organelles that integrate cellular metabolism and apoptotic processes, we assessed mitochondrial function in TECs that were stimulated with kidney injury factors and treated with IL-22 [25]. As shown in **Figure 1A, 1C, 1D**, mitochondrial basal oxygen consumption rate (OCR), maximal respiratory capacity (MRC) and respiration were blocked substantially in the TECs under stress situations. Notably, these abnormalities were largely prevented by IL-22 treatment. Glycolysis in damaged TECs, evidenced by extracellular acidification rate (ECAR), was also markedly restrained, compared with IL-22 plus stimuli-challenged TECs (**Figure 1B, 1E**). Because multiple evidences have suggested that Glut1 plays a principal role in glycolysis and glucose homeostasis [26], we therefore ask whether IL-22 affects Glut1 translocation or expression. As anticipated, we demonstrated that IL-22 not only promoted the translocated Glut1 to the cell membrane surface (PKH26) and Glut1 expression, but also enhanced glucose uptake (**Figure 1F, and 1G**). Overall, these observations indicated that IL-22 could enhance OXPHOS and glycolysis in injured TECs.

**Figure 1.**
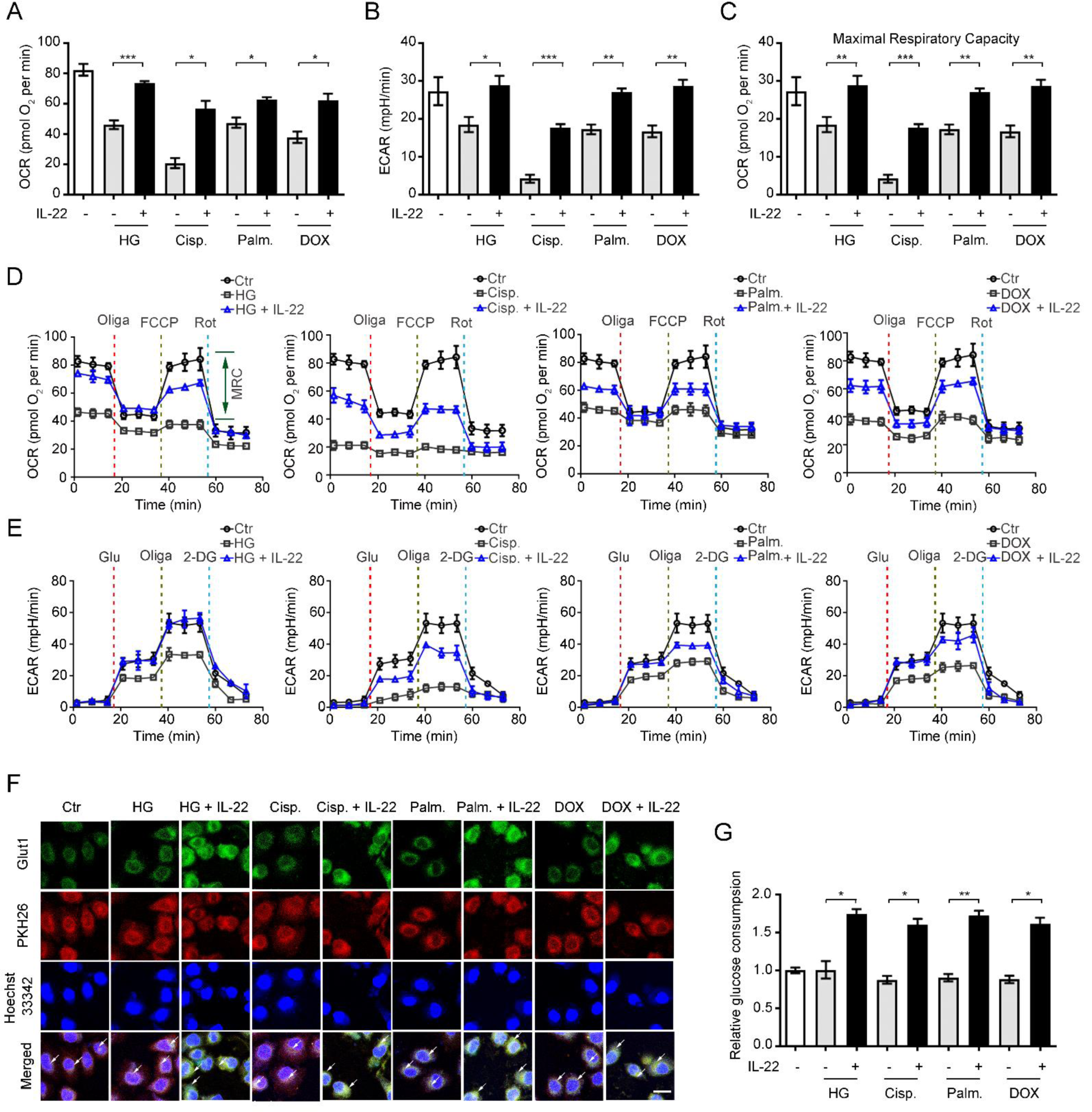
IL-22 preserves mitochondrial fitness and glycolysis in TECs. (**A** and **B)** We use Seahorse XF96 Extracellular Flux Analyzer to test the changes in the extracellular acid rate (ECAR) and oxygen consumption rate (OCR) of TECs. OCR and ECAR in TECs incubated with 50 mM glucose, or 5 μg/mL cisplatin, or 0.2 mM palmitic acid, or 4 μM DOX in the presence or absence of IL-22. (**C**) Maximal respiratory capacity (MRC) of TECs assessed by real-time changes in OCR. (**D** and **E**) Curves in the OCR and ECAR of TECs after incubated with oligomycin, FCCP, glucose, rotenone, and 2-DG. (**F**) Localization and expression of nuclear (blue), Glut1 (green), and plasma membrane (red) in TECs. (**G**) Glucose consumption in TECs after IL-22 treatment for 24 h. *n* = 3; scale bars, 20 μm; Student’s t test (unpaired); ***P < 0.001, **P < 0.01, *P < 0.05.

### IL-22 ameliorates the accumulation of dysfunctional mitochondria in TECs via the activation of mitophagy

We next investigated whether the improvement of cellular metabolism by IL-22 in TECs is attributed to the prevention of mitochondrial dysfunction. As detected by MitoTracker Green and MitoSOX dye, we found that TECs had increased mitochondrial mass and mitochondrial ROS after exposure to stimuli (**Figure 2A and 2B**). To distinguish respiring and dysfunctional mitochondria, the TECs were then stained with MitoTracker Red to detect the mitochondrial membrane potential. Our results revealed that kidney injurious stimuli, such as cisplatin, significantly increased the mitochondria dysfunction (with higher MitoTracker Green and lower MitoTracker Red) (**Figure 2C**). Importantly, IL-22 could preserve mitochondrial fitness in injured TECs, which was showed by the reduction of mitochondrial mass, mitochondrial ROS and mitochondrial dysfunction (**Figure 2A, 2B and 2C**). These observations corresponded to the flow cytometry data and live-cells fluorescence images, wherein the ROS generation and accumulation could be inhibited by IL-22 (**Figure 2D and 2E**). Because mitophagy plays an important role in metabolic processes through restoring metabolic homeostasis and mitochondrial fitness [27], we therefore ask whether IL-22 alleviates mitochondrial dysfunction via induction of mitophagy. Mitophagy was measured by assessing LC3-GFP puncta generation. As expected, IL-22 treatment had increased formation of LC3 puncta in TECs after exposure to stimuli (**Figure 2F**). We then depleted mitophagy to assess its function on the tubule cell metabolism via siRNA-ATG5 (**Figure S1**). Our data suggested that the depletion of mitophagy in TECs significantly inhibited the protective effects of IL-22 in preserving mitochondrial fitness (**Figure 2E**), indicating that IL-22 prevents mitochondrial dysfunctional via the activation of mitophagy.

**Figure 2.**
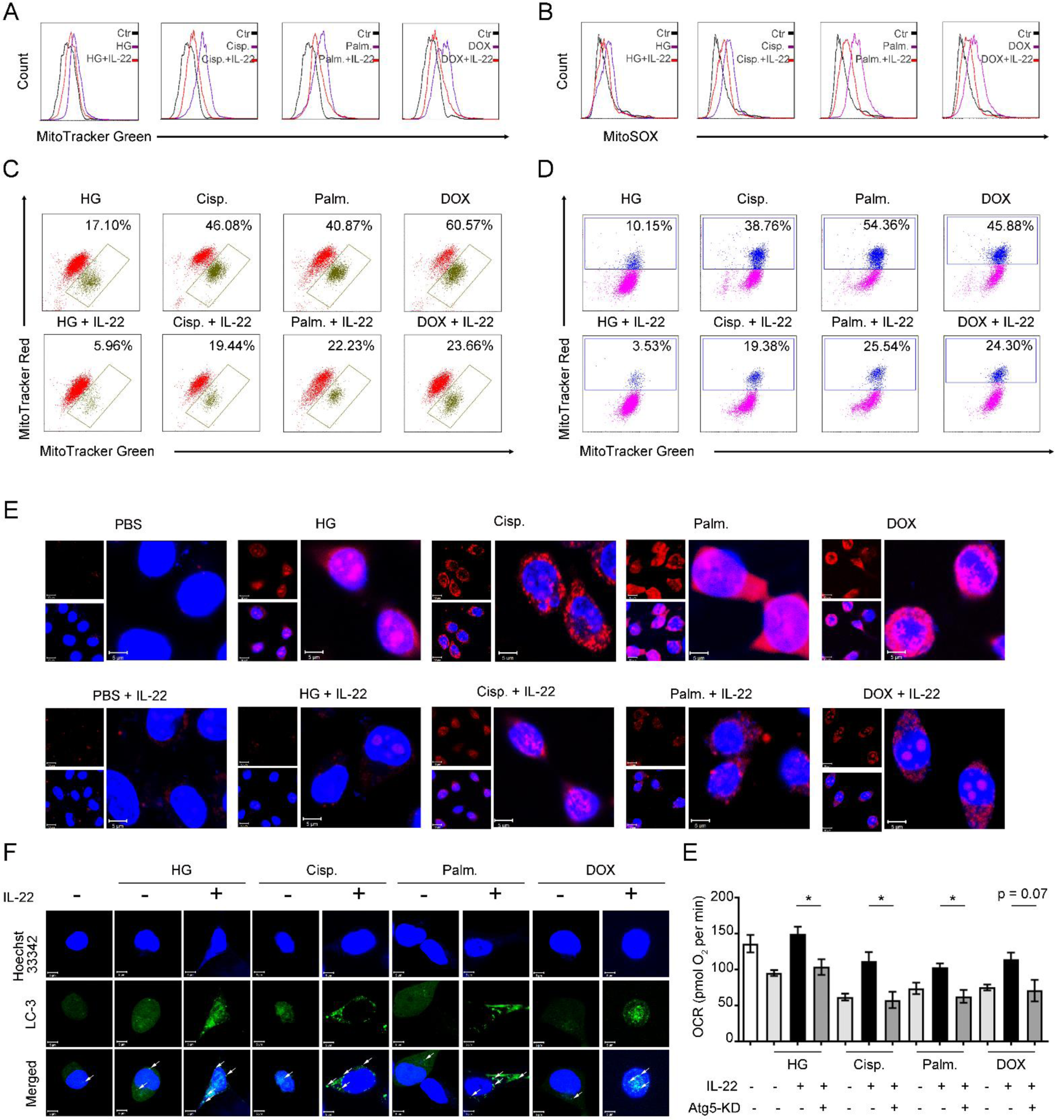
IL-22 inhibits mitochondrial dysfunction and mitochondrial ROS accumulation via the induction of mitophagy. (**A**) Mitochondrial mass was stained and assessed by MitoTracker Green and flow cytometry. Mitochondrial ROS was evaluated in TECs stained with MitoSOX (**B**). Mitochondrial membrane potential was assessed in TECs stained with MitoTracker Green and MitoTracker Red (**C**); mitochondrial ROS accumulation was evaluated by MitoTracker Green and MitoSOX (**D**). (**E**) Confocal image results suggested mitochondrial ROS production labeled with MitoSOX. (**F**) Representative microscopic images indicated LC3-GFP punctate formation in the presence or absence of IL-22 TECs stimulated as (**A**) for 24 hours. (**E**) OCR was assessed in the presence or absence of IL-22 and siRNA-ATG5 for 24 h. *n* = 3; Student’s t test (unpaired); ***P < 0.001, **P < 0.01, *P < 0.05.

### IL-22 maintains mitochondrial integrity of TECs via activation of AMPK/AKT signaling

The AMPK signaling, as a key signaling hub linking cell survival and metabolism, controls glucose metabolism, lipid synthesis and mitochondrial function [28]. According to the renal protective effect of interleukin-22 mediated by metabolic reprogramming of TECs, we assessed whether IL-22 regulates mitochondrial integrity via activation of AMPK signaling transduction. We found that IL-22 increased the activity of STAT3/AMPK/AKT signaling transduction in injured TECs, evidenced by the phosphorylation of STAT3, AMPK, AKT (**Figure 3A**). Additionally, these processes were blocked in tubular cells lacking STAT3, which demonstrated that IL-22 regulated AMPK/AKT signaling through STAT3 (**Figure 3B** and **3C**). We further incubated TECs with Compound C and LY294002 to directly inhibit AMPK/AKT signaling transductions and then evaluate their mitochondrial functions and metabolism states. The results revealed that co-treatment with these signaling inhibitors significantly prevented the improved OXPHOS and glycolytic flux induced by IL-22 treatment (**Figure 3D, 3E, 3F, 3G, 3H** and **3I**), which suggested that IL-22 promoted TECs metabolism via activating AMPK/AKT signaling pathway.

**Figure 3.**
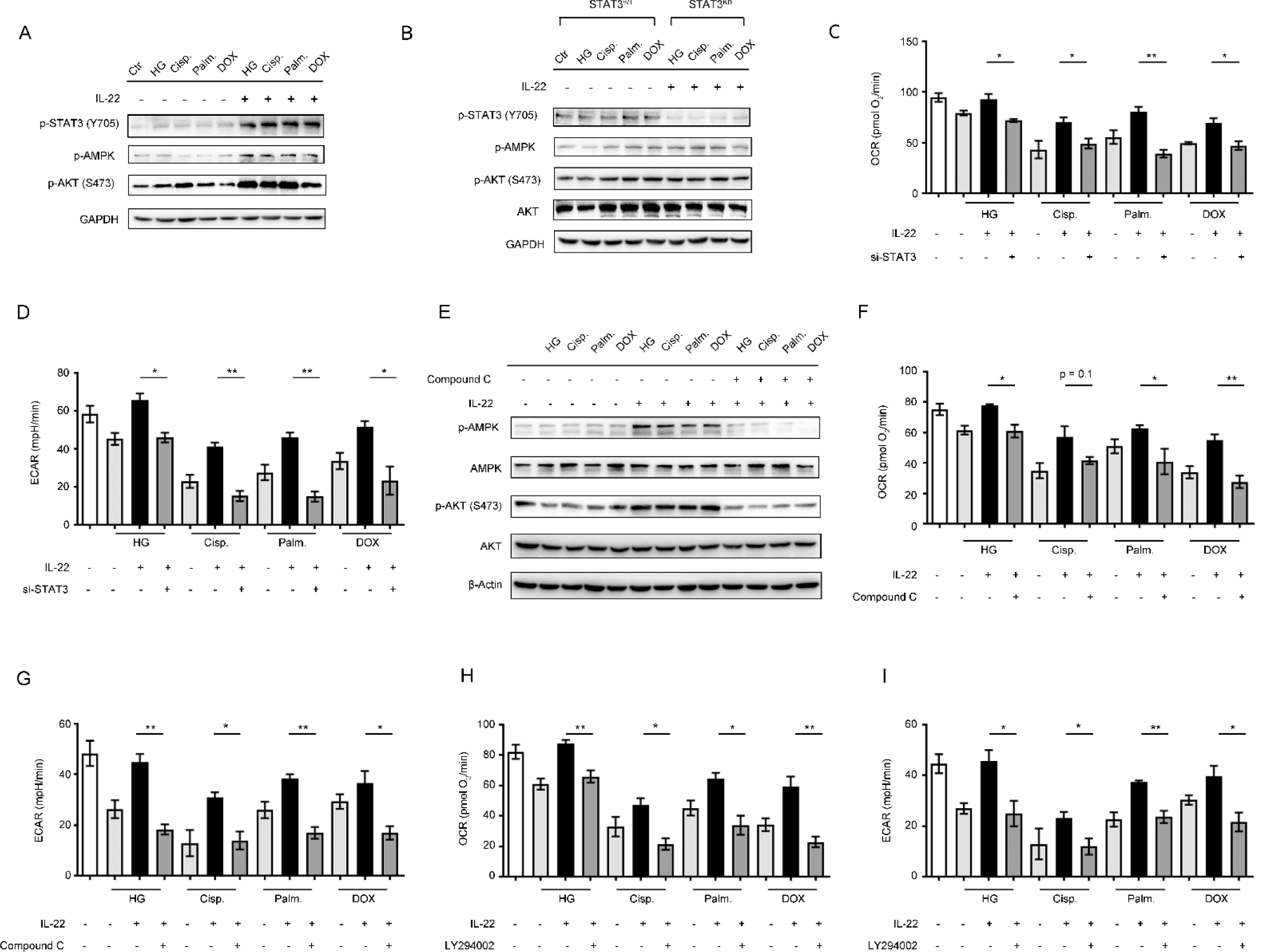
Induction of AMPK/AKT signaling by IL-22 maintains mitochondrial fitness. TECs were stimulated 50 mM glucose, or 5 μg/mL cisplatin, or 0.2 mM palmitic acid, or 4 μM DOX in the presence or absence of IL-22, Compound C, or LY294002 for indicated times. (**A** and **B**) AMPK/AKT signaling pathway activation in TECs or STAT3-WT and STAT3–KD TECs was evaluated by western blot analysis. (**C** and **D**) Changes in the OCR and ECAR of TECs were analyzed. (**E**) AMPK/AKT signaling pathway activation in TECs was evaluated by western blot analysis. (**F, G, H and I**) Changes in the OCR and ECAR of TECs were measured in TECs versus Compound C and LY294002 treated TECs. Student’s t test (unpaired); *n* = 3; ***P < 0.001, **P < 0.01, *P < 0.05.

### IL-22 preserves cellular metabolism and mitochondrial integrity of TECs via induction of PFKFB3

Moreover, to explore how TECs mitochondrial fitness and metabolism were improved, we used RNA sequencing analysis (RNA-seq) to assess gene expression following stimuli injury and IL-22 treatment. We demonstrated that multiple genes were significantly changed in TECs after IL-22 incubation (**Figure 4A**). Using the available databases, the gene set enrichment analysis of those up regulated genes was performed, which suggested a remarkable enrichment of genes in AMPK signaling pathway and MYC targets signaling pathway (**Figure S2**). Notably, PFKFB3 was significantly induced by IL-22 from among the genes during kidney injury factors stimulation (**Figure 4A**). These observations were also indicated by confocal images, Q-PCR and western bolt, and the high expression of PFKFB3 was AMPK/AKT signaling dependent (**Figure 4B, 4C** and **S3**).

**Figure 4.**
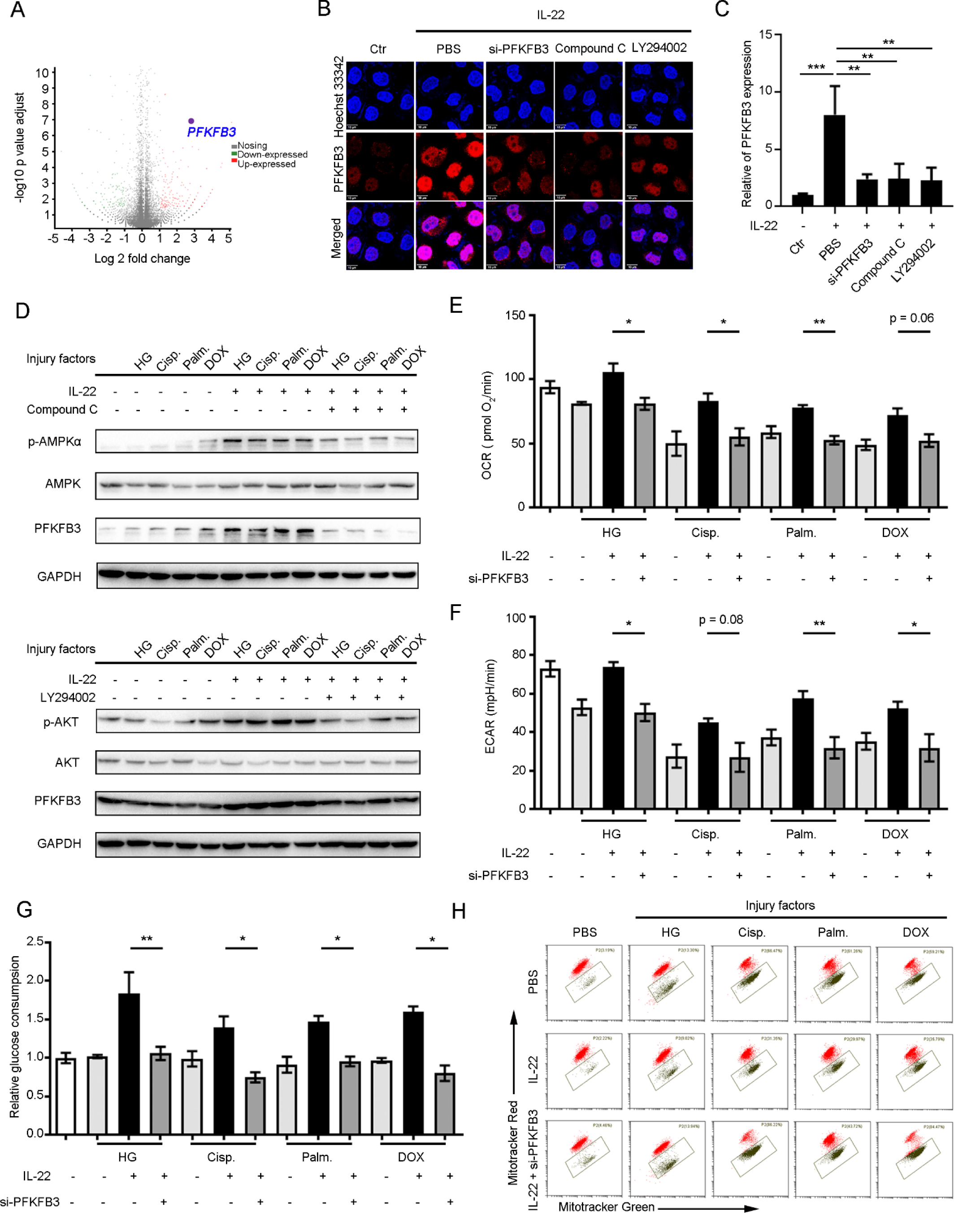
IL-22 maintains mitochondrial integrity and cellular metabolism of TECs via induction of PFKFB3. (**A**) Volcano plot for genes expression in TECs stimulated with glucose, or cisplatin, or palmitic acid, or CCl_4_ in the presence or absence of IL-22 for 24 h. (**B**) Confocal microscopic data demonstrated PFKFB3 overexpression in TECs after IL-22 treatment, which required AMPK/AKT signaling activation. (**C**) Real-time PCR results indicating that IL-22 induced PFKFB3 overexpression in TECs, which could be inhibited by AMPK/AKT signaling blocked. (**D**) The AMPK/AKT/ PFKFB3 signaling pathway activation in TECs after treated with Compound C and LY294002. (**E** and **F**) ECAR and OCR in TECs at the presence or absence of IL-22, or si-PFKFB3 for 24 h. (**G**) Glucose consumption in TECs after IL-22 or si-PFKFB3 treatment for 24 h. (**H**) Mitochondrial dysfunction were evaluated in TECs stained with MitoTracker Red and MitoTracker Green. Student’s t test (unpaired); *n* = 3; ***P < 0.001, **P < 0.01, *P < 0.05.

Previous studies had suggested that PFKFB3 could be regulated by numerous signaling transductions to accelerate epithelial regeneration and repair following injury [29-30]. Thus, we speculated that IL-22 induces PFKFB3 to preserve cellular metabolism of TECs and maintain its mitochondrial fitness. We observed that the inhibition of AMPK/AKT signaling reduced PFKFB3 expression (**Figure 4D**). So, we silenced PFKFB3 to further study its functions in TECs after IL-22 incubation. Our findings indicated that silenced PFKFB3 alleviated the metabolic reprogramming effects of IL-22 on restoring ECAR and OCR (**Figure 4E and 4F**), and prevented glucose uptake as well as dysfunctional mitochondria (**Figure 4G and 4H**). Taken together, these findings indicated that the activation of PFKFB3 plays an important role in mitochondrial fitness and dysfunctional mitochondria elimination after kidney injury.

### IL-22 alleviates kidney injury in cisplatin-induced AKI via suppression renal reactive oxygen species (ROS) accumulation and mitochondrial dysfunction

To test our hypothesis that IL-22 regulates renal metabolic profiles to suppress kidney tubule injury *in vivo*, we initially evaluated renal functions in IL-22 treated mice that had been administered with cisplatin (**Figure 5A**). Four days after cisplatin injection, IL-22 not only substantially reduced tubular cellular damage and hemorrhage; their renal dysfunction also had been significantly attenuated, which were assessed by the serum levels of blood urea nitrogen (BUN) and creatinine (Cr) (**Figure 5B, 5C and 5D**). We further suggested IL-22 prevents kidney injury via enhancing kidney regeneration and activating AMPK/AKT signaling and PFKFB3 (**Figure 5E, 5F and S4**). Next, we employed MitoSOX (mitochondria-specific ROS dye) and JC-1 (mitochondrial membrane potential-dependent dye) to assess the mitochondrial dysfunction involved in cellular metabolism and apoptosis. In the kidney sections of mice that had been treated with cisplatin, there were comparable accumulations of ROS. Of note, at 4 days after IL-22 treatment, the levels of ROS were markedly reduced in the injured kidneys in comparison to ROS from mice after PBS treatment (**Figure 5G**). Taken together, these observations offer evidence that IL-22 plays a protective role in cisplatin-induced AKI via controlling mitochondrial fitness.

**Figure 5.**
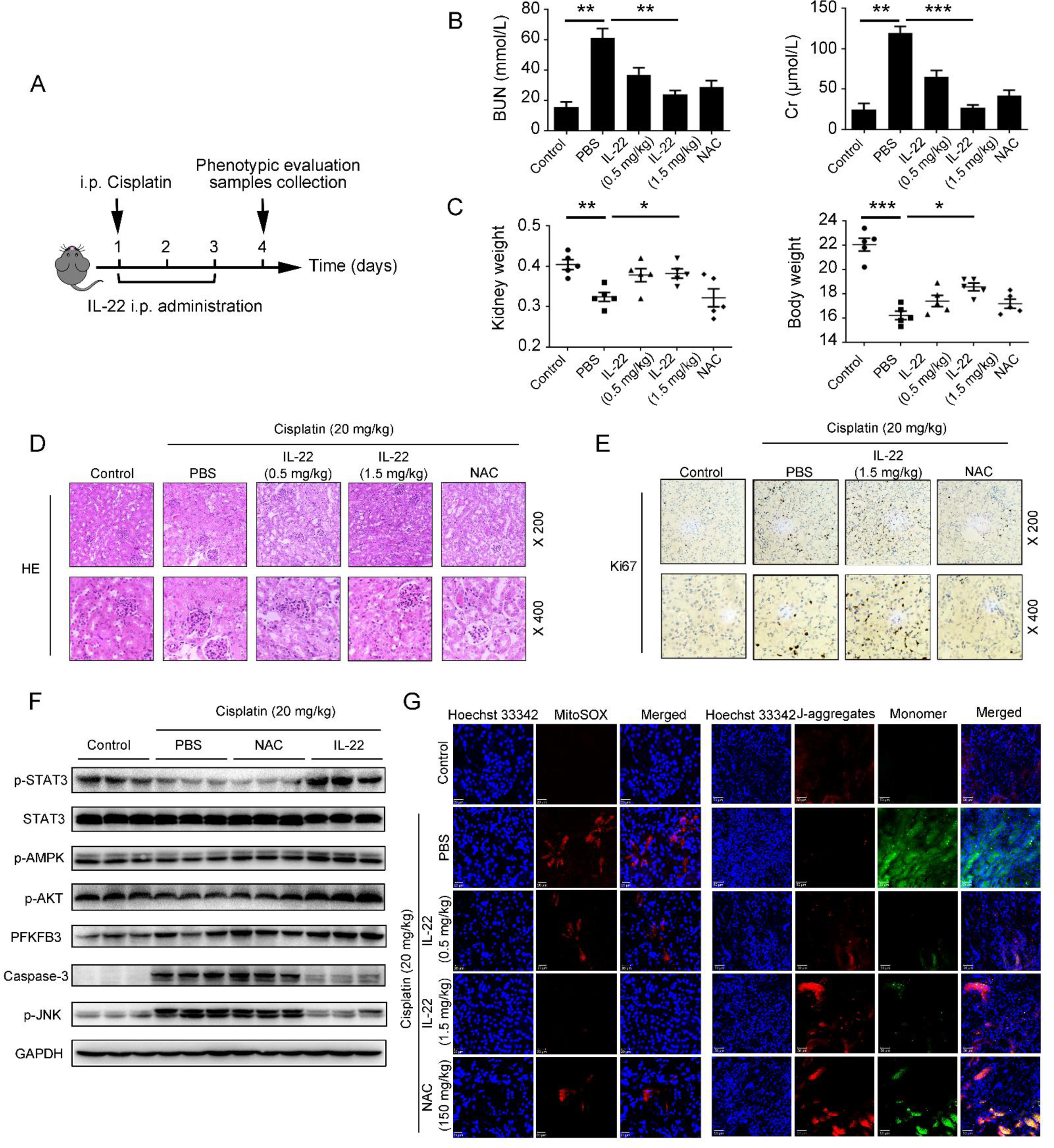
IL-22 alleviates kidney damage in cisplatin-induced AKI via inhibition renal ROS accumulation and dysfunctional mitochondria. (**A**) Schematic diagram of the experimental protocols to evaluate the protective effects of IL-22 in cisplatin (20 mg/kg) induced kidney damage. PBS as a vehicle control group and *N*-acetyl-L-cysteine (NAC, 150 mg/kg) as a positive control group. (**B** and **C**) Serum BUN levels, serum Cr levels, kidney weights and body weights were assessed. Representative HE (**D**), Ki-67 staining (**E**), MitoSOX (**G**) and JC-1 (**G**) images of the kidney sections were presented. (**F**) Comparison of STAT3/AMPK/AKT/PFKFB3 activation in kidney extracts from IL-22-treated mice was evaluated by western blot.

### IL-22 ameliorates diabetes-induced renal injury via inhibition of mitochondrial dysfunction through the activation of AMPK/AKT signaling and PFKFB3

We and others have demonstrated that IL-22 could exert protective potency in diabetic nephropathy (DN), which is the main cause of ESTD [31-32]. To further study the underlying mechanisms, we fed db/db mice with high-fat diets (HFD) and then treated IL-22 (2.5 mg/kg, ip) for 8 weeks (**Figure 6A**). When mice were fed with HFD only, we found kidney tubular fibrosis and dilatation, in addition to atrophy. In contrast, when mice were treated with IL-22, renal fibrosis and histopathology were significantly improved, as well as kidney functions. Enhanced kidney functions were also demonstrated by decreased serum Cr, BUN levels and total urinary albumin/24 h (UAE) levels (**Figure 6B, 6C, 6D and 6E**). To study whether these activities are associated with mitochondria, we assessed the mitochondrial function involved in cellular metabolism and apoptosis. The results indicated that the increased mitochondrial ROS levels and accumulated dysfunctional mitochondria were significantly alleviated by IL-22 treatment (**Figure 6F and 6G**). To further study whether the protective functions of IL-22 in kidney are mediated through PFKFB3, we injected mice with PFKFB3 shRNA adenovirus to knockdown PFKFB3. Our data indicated that the beneficial effects of IL-22 on HFD-induced renal injury, necrosis, steatosis, mitochondrial dysfunction and ROS accumulation and the activation of related signaling pathways were significantly blocked by PFKFB3 knockdown (**Figure 6B, 6C, 6D, 6E, 6F and 6G**). Additionally, the increased expression of AMPK/AKT signaling was also observed after treatment with IL-22 (**Figure 6H**). Along with the alleviation of kidney dysfunction, the expression of PFKFB3 and important enzymes in cellular metabolism were also improved in kidney of IL-22-treated mice (**Figure 6H** and **6I**). Overall, our data suggested that IL-22 alleviated kidney damage via activation of AMPK/AKT signaling transduction, inhibition of ROS accumulation and mitochondrial function regulation through PFKFB3.

**Figure 6.**
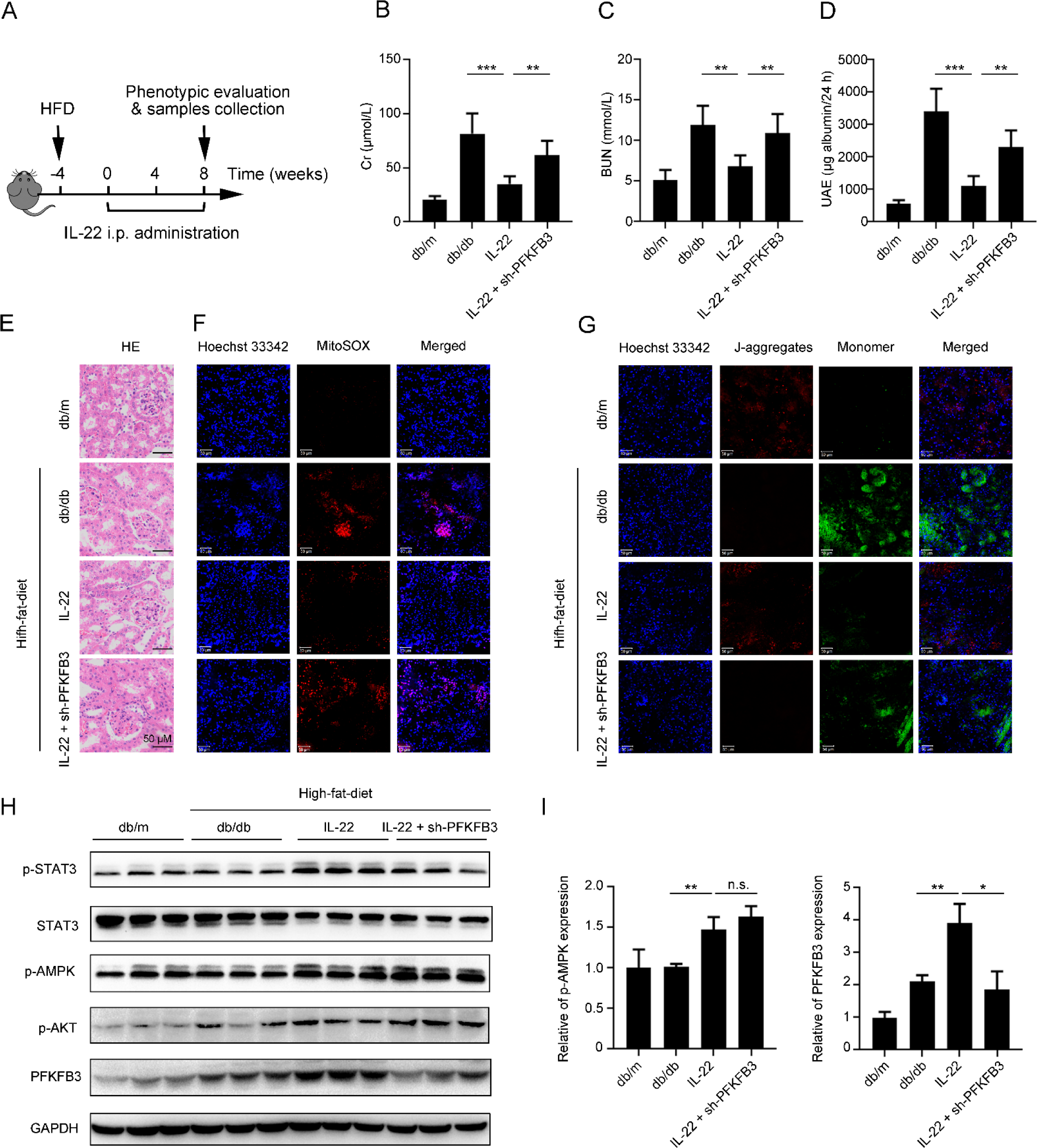
IL-22 alleviates diabetes-induced renal damage through AMPK/AKT signaling and inhibition of dysfunctional mitochondria via activation of PFKFB3. (**A**) Schematic diagram of the experimental protocols to evaluate the protective effects of IL-22 in diabetes-induced renal damage. Db/m as a vehicle control group. (**B, C** and **D**) Serum BUN levels, serum Cr levels, and UAE were evaluated. Representative HE (**E**), MitoSOX (**F**) and JC-1 (**G**) images of the kidney sections were presented. (**F**) Comparison of STAT3/AMPK/AKT/PFKFB3 activation in kidney extracts from IL-22-treated mice was evaluated by western blot.

## Discussion

IL-22 is secreted by immune cells, such as NK cells, neutrophils, innate lymphoid cells (ILCs) and T cells, but does not directly affect these cells. It predominantly regulates the functions of epithelial cells owing to the restricted IL-22R1 expression on epithelial cells including TECs [15, 33]. Previously studies have suggested that IL-22 can protect against renal epithelial injury and accelerate tubular regeneration through inhibition of NLRP3 inflammasome and activating STAT3 and AKT signaling [16, 31]. However, the underlying mechanisms of this protection are not yet to be elucidated. To further determinate the molecular basis of IL-22’s function in kidney injury, we first investigate whether IL-22 can alleviate kidney cell dysfunction and subsequent apoptosis through regulating their metabolic states. Moreover, we studied two models of kidney injury diseases (AKI and DN) with IL-22 administration and compared the observations with those of control groups. We indicated that IL-22 corrects metabolic reprogramming to maintain mitochondrial integrity, decreases mitochondrial ROS accumulation in injured TECs, and alleviates progressive kidney damage and necrosis. Moreover, our findings suggested that IL-22 could activate AMPK/AKT associated signaling pathways in TECs and kidney, important mediators of epithelial wound healing and cell survival, to ameliorate mitochondrial dysfunction and the deteriorating metabolic profiles and opening up a novel field for IL-22 mediated renal protective mechanism. Overall, our results indicated that the renal protective effects of IL-22 were mediated via metabolic reprogramming processes.

The critical role of OXPHOS and glycolysis in kidney protection have been revealed in previous studies, where balancing fuel utilization by inhibitory S-nitrosylation of pyruvate kinase M2 (PKM2) protects against kidney injury [19]. Our results suggesting the promotion of OXPHOS and glycolysis by IL-22 via increasing of metabolic regulators expression and glucose uptake demonstrate that IL-22 reverses the deteriorating metabolic states associated with the kidney injury. These observations are consistent with the renal protective functions of recombinant irisin and also in line with findings from hepatocytes, where both OXPHOS and glycolysis is needed for cellular functions, but if these are inhibited, then the primary tubule cells and hepatocytes become dysfunction [34-35].

Mitochondrial dysfunction has emerged as key molecular basis that integrate metabolic profiles and cell death. Herein, we illustrate that upon injury factors stimulation, TECs treatment with IL-22 had ameliorated accumulation of mitochondrial ROS and dysfunctional mitochondria as compared with the decreased mitochondrial integrity and function in control groups. Importantly, AMPK signaling pathways, critical energy sensors, modulate metabolic processes to maintain cellular homeostasis and prevents cell or tissue damage [36-37]. In current study, our observations indicating that IL-22 preserves mitochondrial integrity and function via activating the AMPK/AKT signaling demonstrate that IL-22 prevents mitochondrial dysfunction in a direct method, which is vital to maintaining TECs respiratory capacity. Furthermore, we demonstrated that IL-22 signaling through STAT3 has direct effects on preserving mitochondrial fitness, as those have been investigated previously that induced STAT3 phosphorylation is critical for preserving mitochondrial integrity [38].

Multiple literatures have previously indicated PFKFB3 plays the part of central modulator in cell metabolism [29-30, 40]. Herein, we first demonstrate that IL-22 promotes PFKFB3 activation through AMPK/AKT signaling in TECs, suggesting IL-22 regulates metabolic states via involving control of PFKFB3. Moreover, previous studies further indicated that PFKFB3 is important to control the exaggerated OXPHOS and glycolysis in lots of cells or tissues [41-42]. Consistent with these studies, our observations also indicate a promotion of OXPHOS and glycolysis in IL-22-treated TECs via the activation of PFKFB3. Among numerous downstream modulators of IL-22 have been investigated, our data illustrate that PFKFB3 is significantly upregulated after IL-22 treatment during TECs damage and that the STAT3–AMPK/AKT-PFKFB3 axis is vital for the preservation of mitochondrial integrity during kidney injury. They remain to be investigated if IL-22 also regulates other metabolic regulators and processes related with kidney injury through activating the STAT3–AMPK/AKT-PFKFB3 signaling pathway.

## Conclusions

In conclusion, we demonstrated the therapeutic potential and underlying mechanisms of IL-22 in kidney damage. IL-22 regulated renal cell metabolism via a metabolic reprogramming to preserve mitochondrial integrity. Inhibition of their regulators (i.e., AMPK, AKT, PFKFB3) could lead to abnormal metabolic profiles and loss of mitochondrial fitness, as indicated in TECs with spontaneous IL-22 treatment, which had increased dysfunctional mitochondria and mitochondrial ROS accumulation. Most importantly, the kidney damage by mitochondrial dysfunction and ROS accumulation in mice models was substantially alleviated by IL-22 treatment through the activation of STAT3-AMPK-AKT-PFKFB3 signaling. Thus, our study indicated that targeting associated metabolic signaling could be directly favorable for numerous kidney damage diseases and IL-22 is a potential therapeutic agent for preventing and treating these diseases.

## Materials and methods

### Reagents

Oligomycin, rotenone and cyanide p-trifluoromethoxyphenyl-hydrazone (FCCP) were obtained from Seahorse Biosciences, USA; recombinant IL-22 was provided by Novoprotein, China; cisplatin (Cisp.), MitoTracker Green, LY294002, 5,5′,6,6′-tetrachloro-1,1′,3,3′-tetraethylbenzimidazolylcarbocyanine iodide (JC-1), Compound C were obtained from Beyotime Biotechnology, China; antibodies targeting Glu1, β-Actin, p-AMPKα, AMPK and GAPDH were purchased from Abcam, USA; antibodies for STAT3, p-STAT3 (Y705), AKT, p-AKT (S473), PFKFB3, Capase-3 were purchased from Cell Signaling Technology, USA; rapamycin, palmitic acid (Palm.), glucose were obtained from Sigma-Aldrich, USA; MitoSOX, Hoechst33342, MitoTracker Red, PKH26 were obtained from Invitrogen, USA.

### Cell culture

Human proximal tubule epithelial cell (HK-2) was obtained from Cell Bank of Shanghai Institute of Biochemistry and Cell Biology, Chinese Academy of Sciences and maintained following standard protocols. HK-2 was seeded in 6 well plates (1 × 10^6^ cells/well) or 96 well plates (1 × 10^5^ cells/well) and cultured in 90% RPMI1640 medium (Gibco, USA) supplemented with 100 U/mL penicillin, 100 μg/mL streptomycin and 10% fetal bovine serum (FBS) (Gibco, USA). We first incubated HK-2 with IL-22 (0.5 μg/mL) for 0.5 h, then 50 mM D-glucose or 5 μg/mL cisplatin or 0.2 mM palmitic acid or or 4 μM doxorubicin (DOX) for 24 h. In some cases, cells were cultured in the presence of Compound C (5 μM), LY294002 (20 μM) or rapamycin (50 nM) for indicated times.

### Seahorse experiments

We analyzed TECs using a Seahorse XF96 Extracellular-Flux-Analyzer for the changes in the extracellular acid rate (ECAR) and oxygen consumption rate (OCR) and. Briefly, HK-2 (1 × 10^5^) was grown on 96-well Seahorse-plates for overnight. Then, we washed cells with seahorse assay medium containing 1 mM pyruvate, 10 mM glucose and 2 mM glutamine (or glucose free for testing ECAR) in an atmosphere without CO_2_ at 37°C for 0.5 hour. Experimental data were recorded after the under-mentioned inhibitors were injected at an optimum concentration of FCCP (1.0 µM), oligomycin (1.0 µM), rotenone/antimycin A (0.5 μM), 2-DG (50 mM) or glucose (10 mM).

### Animal models

Male BALB/c mice (6-8 weeks) were purchased from Slaccas Experimental Animal Co. (Shanghai, China). Db/db and Db/m mice were provided by Model Animal Research Center of Nanjing University (Nanjing, China). All mice were kept in specific pathogen free (SPF) facilities and grown following standard protocols. For the AKI model, BALB/c mice received cisplatin (20 mg/kg) or saline control by intraperitoneal injection. Db/db mice were fed with high-fat-diet (HFD) at the indicated time points to induce diabetic nephropathy (DN). Db/m mice were fed with chow-diets as control group. Our animal experimental procedures were carried out following the protocols evaluated and approved by the Animal Care and Use Ethics Committee of School of Pharmacy, Fudan University. Immunohistochemical staining, measurement of mitochondrial membrane potential, histological and ROS staining of kidney sections were performed as previously described [22-24].

### Real-time PCR

We obtained the total-RNA by TRNzol reagents (Beyotime Biotechnology, China) from cell or tissue samples and then transcribed to cDNA using a MMLV-reverse transcriptase test kit (Beyotime Biotechnology, China). The expression level of mRNA was analyzed by the real-time PCR instruments (BioRad, USA) using SYBR green qPCR-mix assay kit (Beyotime Biotechnology, China) and standardized to GAPDH.

### Flow cytometry

Kidney cells were grown and treated as described above. MitoSOX (mitochondrial ROS), MitoTracker Green (total mitochondrial mass) and MitoTracker Red (mitochondrial membrane potential) staining were carried out according to the manufacturer’s instructions and previous researches [22-24]. We obtained the results by the Beckman Coulter Flow Cytometer and the CytExpert software (BD Biosciences, USA).

### Glucose uptake

We measured the glucose concentrations in cell culture supernatant by a glucose uptake assay test according to the manufacturer’s protocol and previous researches manufacturer’s (GAHK20, Sigma, USA). Results normalization was carried out according to the number of cells.

### Gene knockdown

We used the small interfering RNA (siRNA, RiboBio, China) to knockdown specific gene. Firstly, Lipofectamine RNAiMAX and siRNA (100 pmol) were gently mixed via pipetting and incubated at 37°C for 0.5 hour. Then we changed the standard cell culture medium to transfection cocktail and incubated the cells for 6 hours for siRNA gene transfection. Next, we changed the transfection cocktail to fresh cell culture medium. After 48 hours, kidney cells were incubated with IL-22 for further studies.

### Western blots

We extracted total proteins from cell or tissue by RIPA lysis buffer (Beyotime Biotechnology, China). Then, we collected the lysates by centrifugation at 4 °C at 12000 rpm for 10 min. Subsequently, the lysates were heated with the sample buffers at 100 °C for 15 min, subjected to sodium dodecyl sulfate polyacrylamide gel-electrophoresis (SDS–PAGE) and electro transferred to polyvinylidene difluoride (PVDF) membrane (Millipore, Germany). The membranes were then blocked with 5% bovine serum albumin (BSA) for 2 hours and co-incubated with primary antibodies at 4°C overnight. After washing three times, the membranes were incubated with horseradish peroxidase-conjugated secondary antibodies (CST, USA). The protein blots were visualized by the enhanced chemiluminescence imaging system.

### Immunofluorescence

The kidney cells were fixed by 3%-4% paraformaldehyde and permeabilized with 0.1% Triton X-100. Then, we blocked the cells with 5% bovine serum albumin (BSA) for 2 hours. Cells were incubated overnight with anti-GLUT1 antibody at 4°C, washed four times, and then stained with PKH26. Subsequently, kidney cells were stained with Hoechst 33342 and mounted on slides. Data were obtained by the confocal microscopy.

### Statistics

Data were expressed as means ± standard deviations (SD) and were shown using GraphPad Prism 5.0. For results comparing two or more groups, differences were calculated using the Student’s t-test or the one-way of analysis (ANOVA). ***, **, and * suggested P < 0.001, P < 0.01 and P < 0.1, respectively.

## Acknowledgments

This study was funded by the China Postdoctoral International Exchange Program and National Natural Science Foundation of China (No. 81773620, 31872746).

---

**Figure S1.**
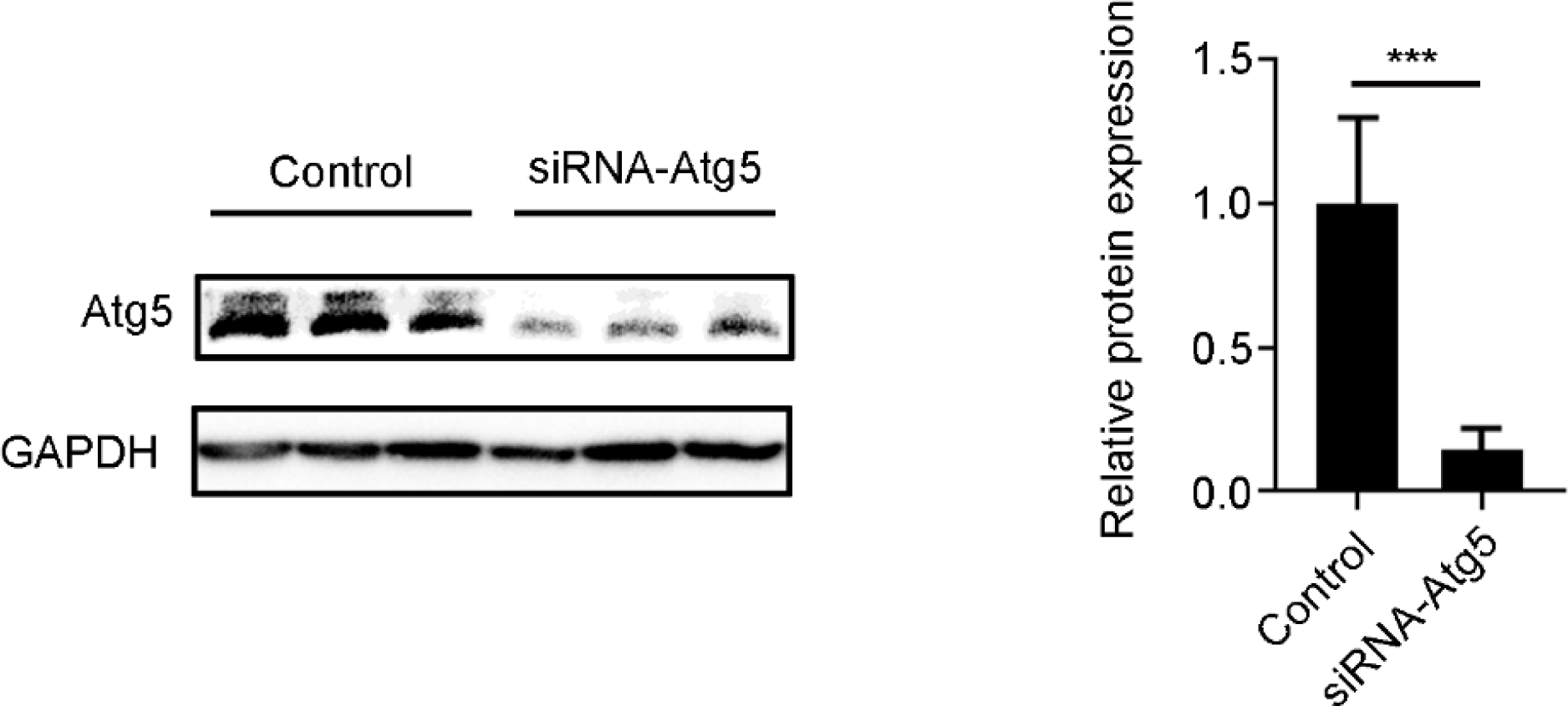
TECs were treated with PBS or siRNA-Atg5 for 48 h. Atg5 expression in TECs was evaluated by western blot analysis. Densitometric values were quantified and normalized to control group (*n* = 3; mean ± SD; \*\*\**P* < 0.001). The values of control group were set to 1.

**Figure S2.**
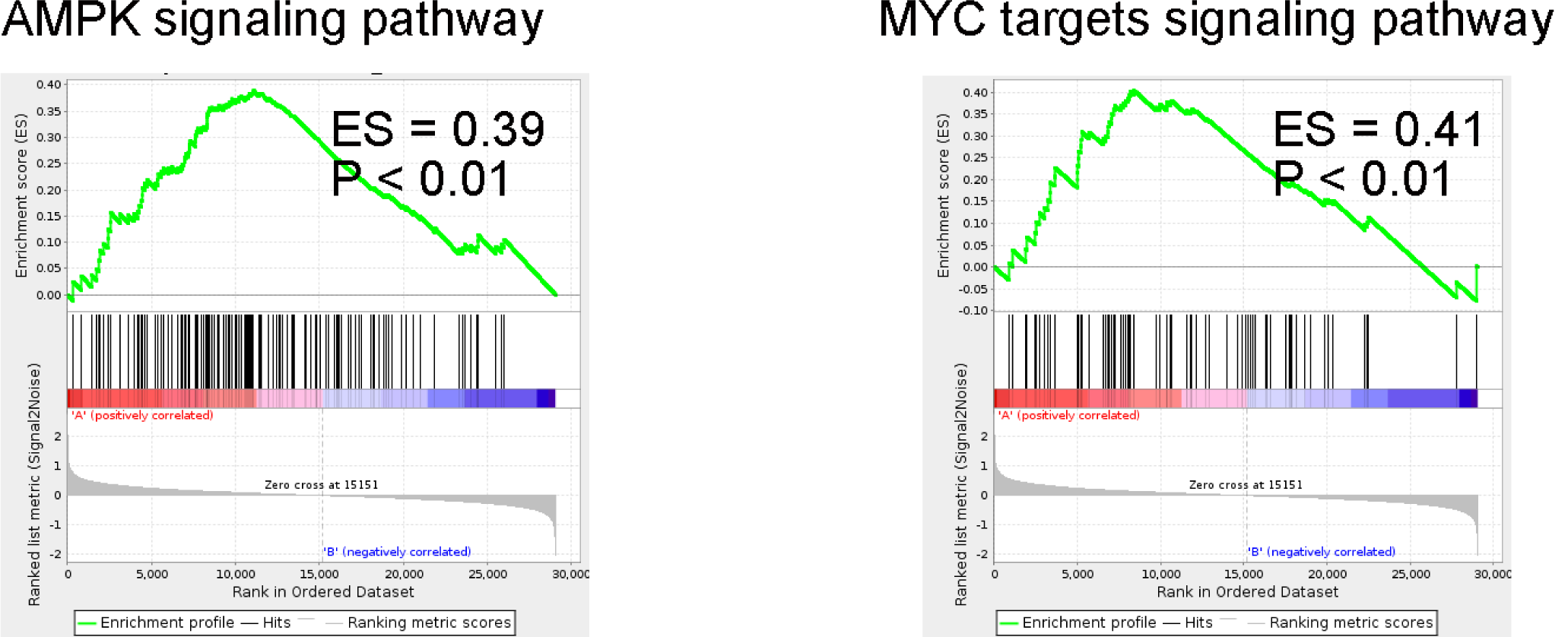
KEGG of AMPK signaling pathway and MYC targets signaling pathway in IL-22-protected and -nonprotected TECs.

**Figure S3.**
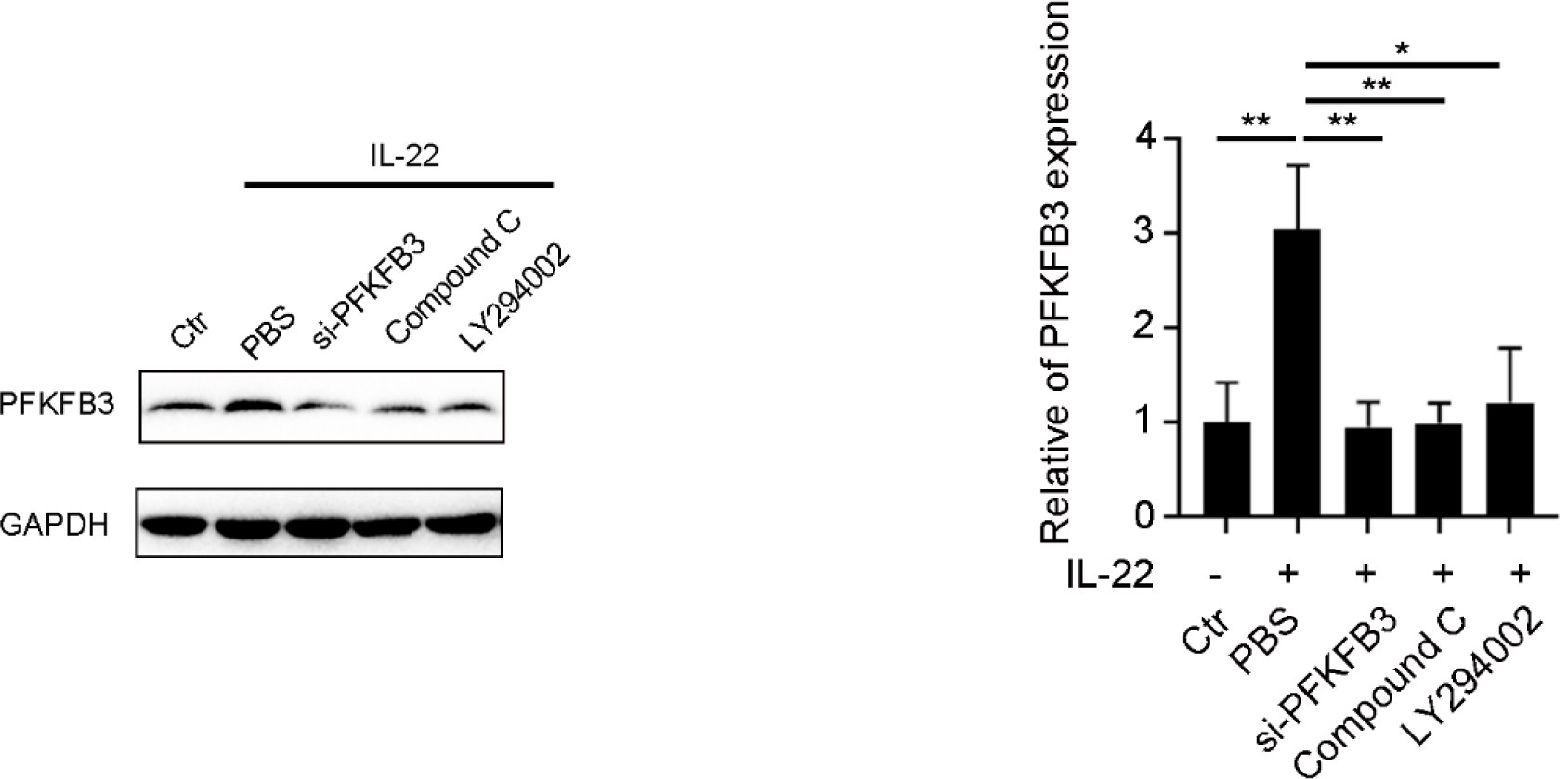
Kidney cells were treated with PBS or siRNA-PFKFB3, or Compound C, or LY294002 or IL-22. PFKFB3 expression was evaluated by western blot analysis. Densitometric values were quantified and normalized to control group (*n* = 3; mean ± SD; \*\*\**P* < 0.001). The values of control group were set to 1.

**Figure S4.**
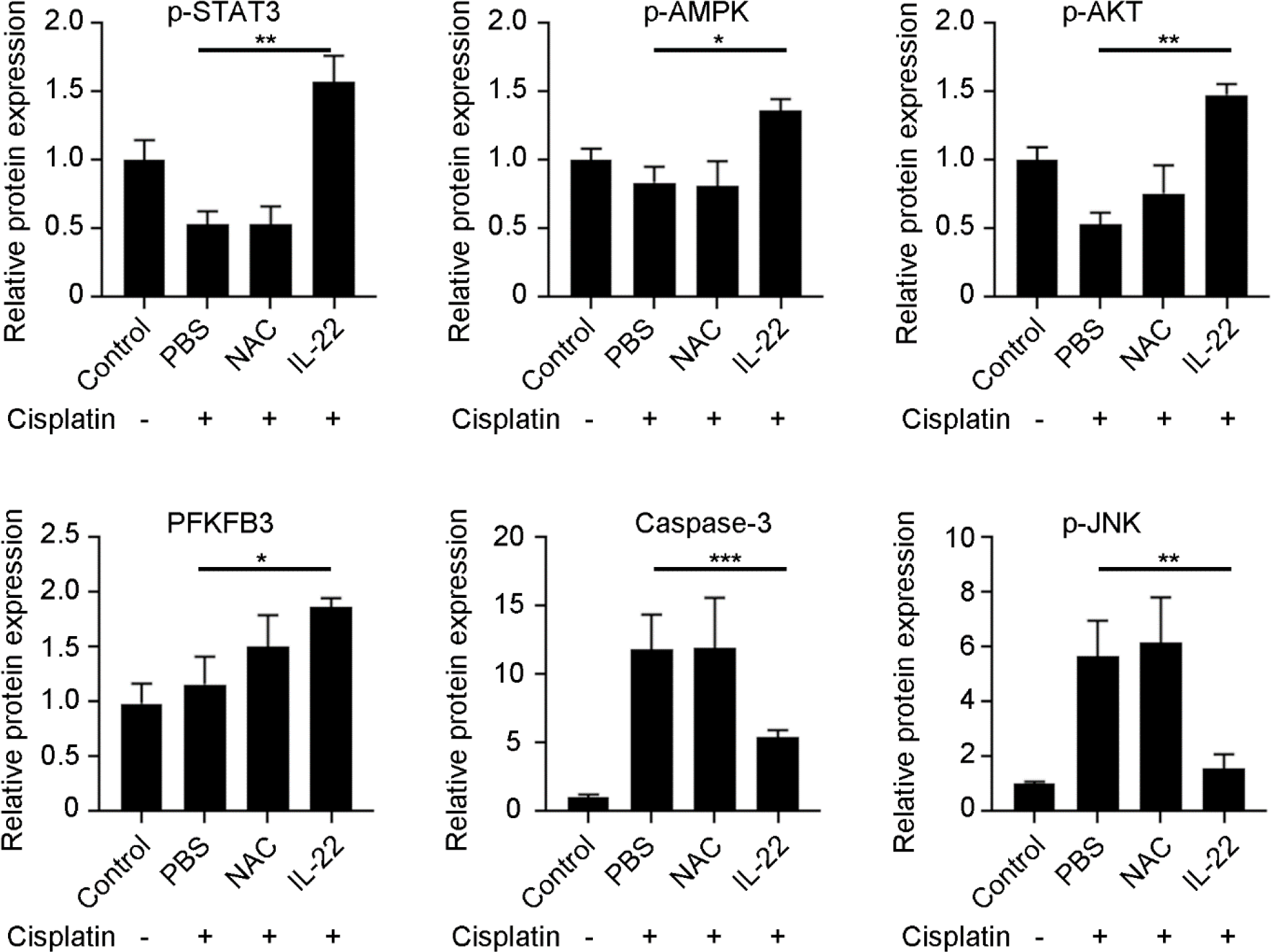
Densitometric values were quantified and normalized to control group (*n* = 3; mean ± SD; \**P* < 0.05). The values of control group were set to 1.

